## Supplementary Information for "CRISPR-based gene drives generate super-Mendelian inheritance in the disease vector *Culex quinquefasciatus*"

### SUPPLEMENTARY FIGURES

#### Supplementary Figure 1

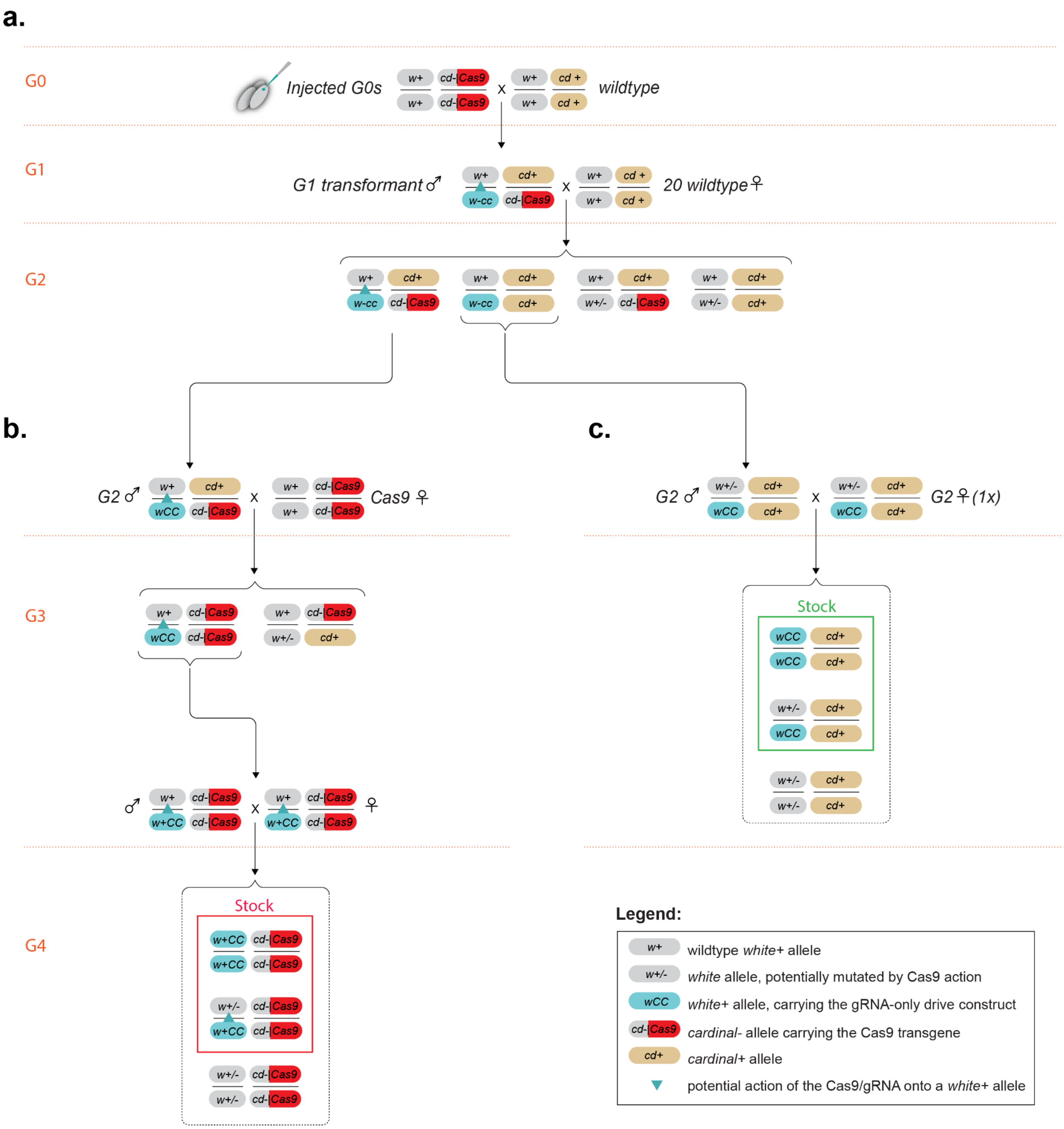


**Supplementary Figure 1. Cross schemes used to isolate two separate *Culex quinquefasciatus* lines carrying the *white*-gRNA6 homing element. (a)** Depicts the genetic crosses performed to generate G1 transformants; the only G1 transformant isolated was male and was crossed to ~20 wild-type females to obtain G2 offspring. **(b)** Several G2 males positive for both transgenes (*vasa*-Cas9 and *white*-gRNA6) were crossed to virgin females homozygous for the *vasa*-Cas9 line, to obtain G3 offspring, which were intercrossed to generate G4 offspring from which only mosquitoes positive for both transgenes were selected to establish a dual-transgene line. **(c)** G2 males and the single G2 female recovered carrying only the eGFP transgene were intercrossed to obtain G3 offspring; from these individuals, only eGFP+ mosquitoes were selected to establish a line carrying only the *white*-gRNA6 transgene.

### Supplementary Figure 2

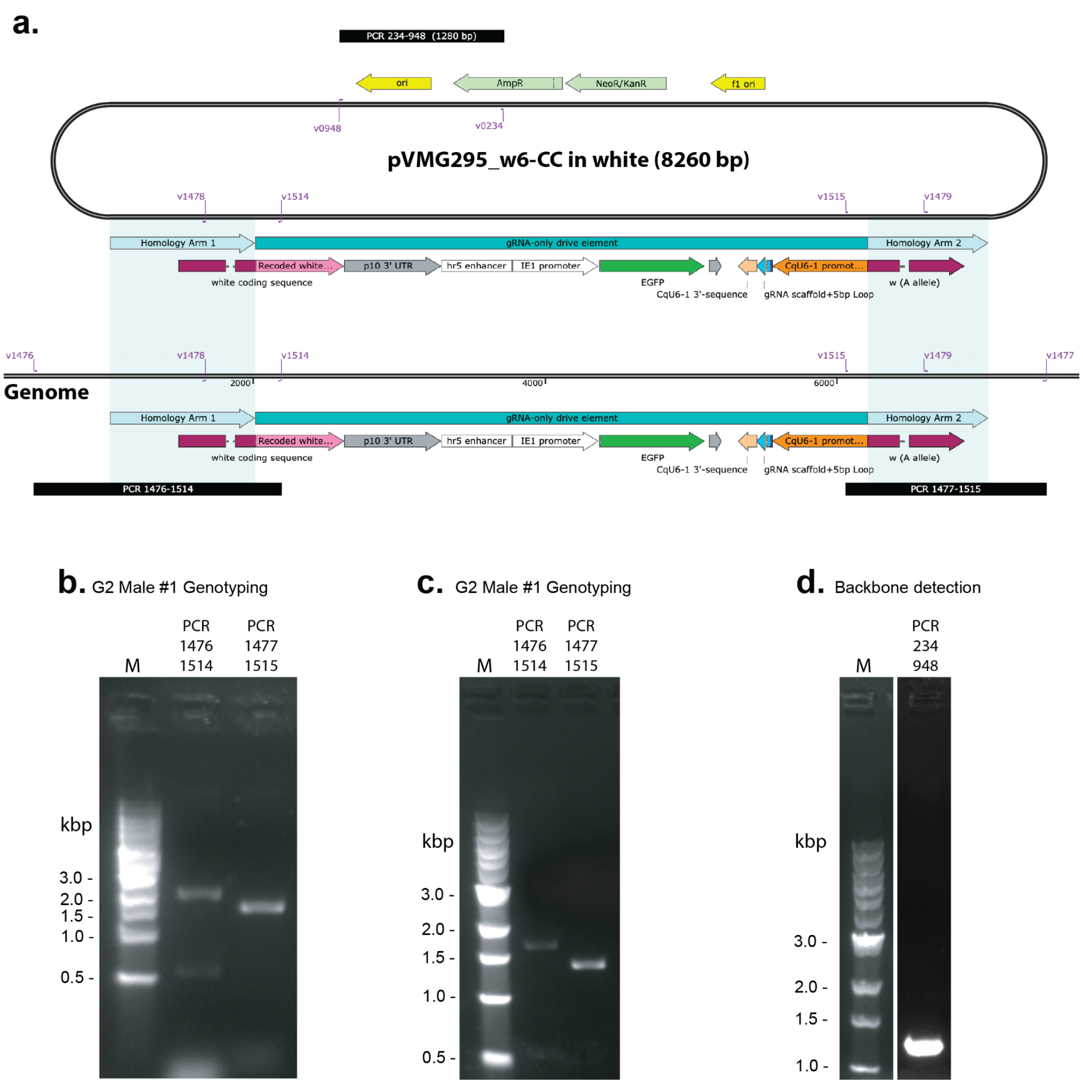


**Supplementary Figure 2. Molecular characterization of the *white*-gRNA6 drive transgenic insertion. (a)** Schematic of the transgene inserted in the genome, along with the plasmid construct used for the transgenesis; teal shading highlights the homology arms used for the transgene insertion by HDR; highlighted in purple are the locations of the primers used for the molecular characterization; black bars indicate the position of the PCR amplicons used for genotyping. **(b)** Gel image of genotyping PCR performed on an eGFP+ G2 male (see Supplementary Figure 1a); **(c)** Gel image of genotyping PCR performed on an eGFP+ G2 female (see Supplementary Figure 1a); **(d)** Gel image of genotyping PCR performed on an eGFP+ male and detecting the presence of the backbone in the transgene, suggesting the presence of tandem insertions.

**Supplementary Figure 3**

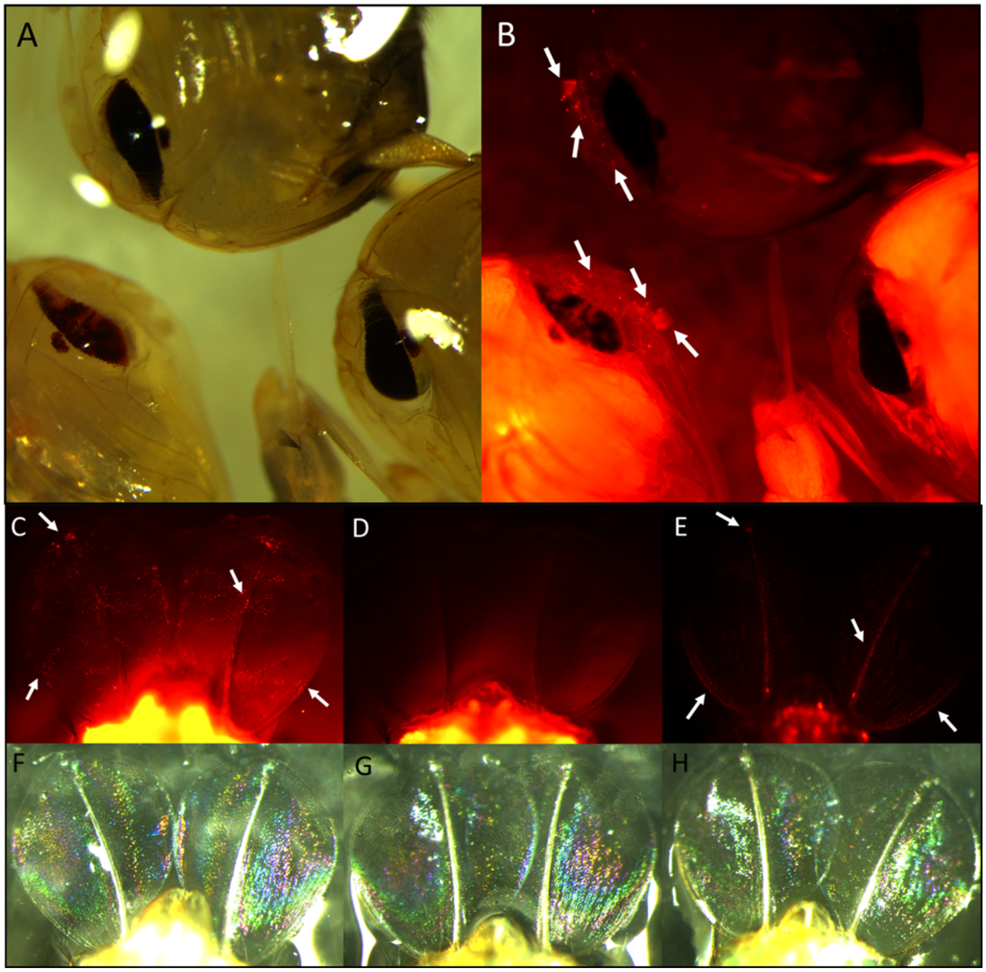


**Supplementary Figure 3: Visual identification of vasa-Cas9 x kmo-gRNA transheterozygotes. (A-B)**: Image of kmo-gRNA heterozygote (upper individual), vasa-Cas9 x kmo-gRNA double heterozygote (lower left individual) and vasa-Cas9 heterozygote (lower right individual). **(A)** taken under white light, **(B)** under mCherry filter. Double heterozygotes could be distinguished from vasa-Cas9 heterozygotes by the presence of 'punctate' nuclear localized DsRed fluorophore signals in the 'forehead' region of pupae which houses the base of the developing proboscis (white arrows). These were also present in the kmo-gRNA heterozygote, however this line lacked the full-body DsRed fluorescence of the vasa-Cas9 insertion. **(C)**, **(F)**: vasa-Cas9 x kmo-gRNA double heterozygote. **(D)**, **(G)**: vasa-Cas9 heterozygote. **(E)**, **(H)**: kmo-gRNA heterozygote. A similar pattern of punctate DsRed fluorescence could be observed in the paddles of these pupae—appearing in the double heterozygote and the kmo-gRNA heterozygote (white arrows), but not in the vasa-Cas9 heterozygote. **(C)**, **(D)** and **(E)** taken under mCherry filter. **(F-H)** White light photographies. Taken together, punctate fluorescence in these two regions, alongside full body fluorescence, was used to discern double heterozygotes from other genotypes.

**Supplementary Figure 4**

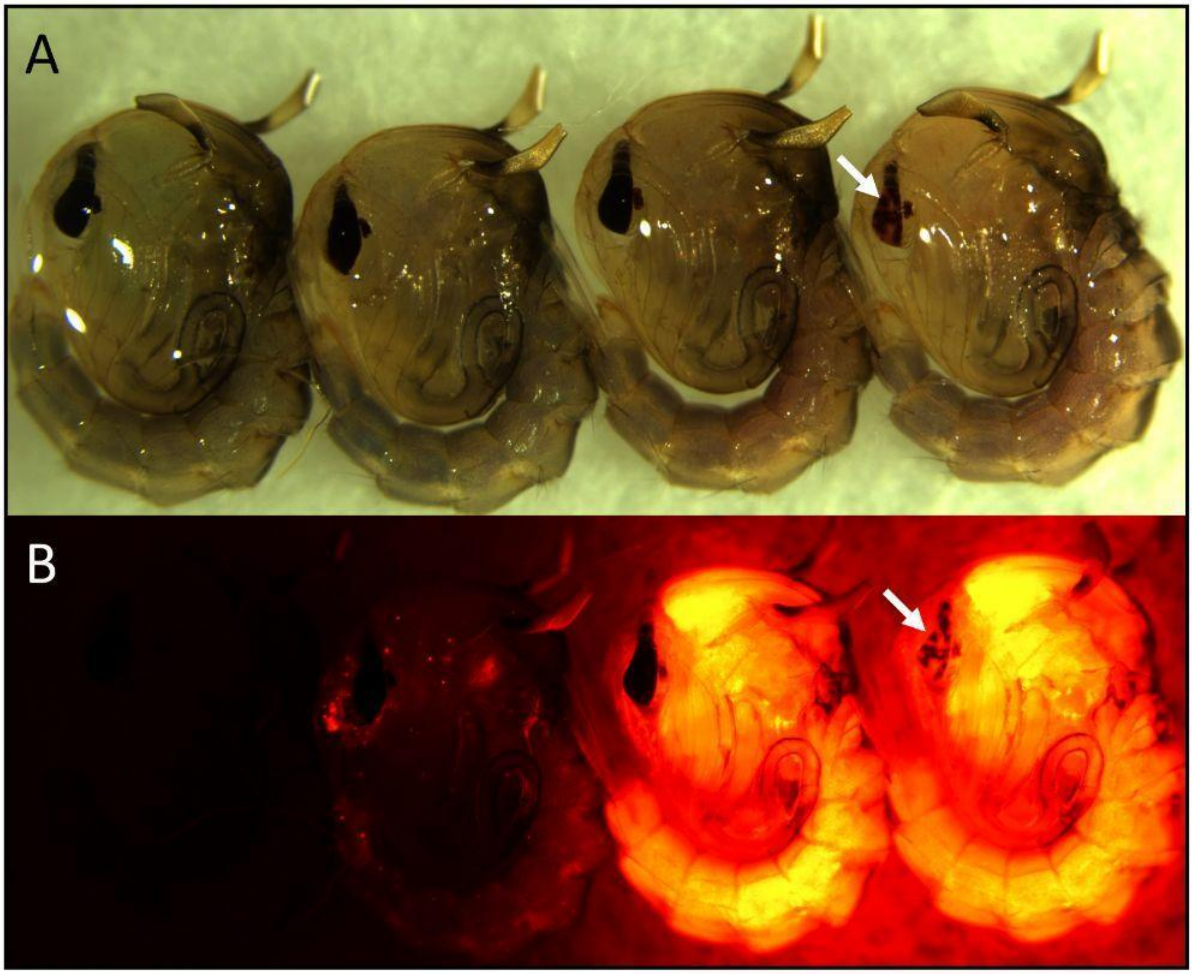


**Supplementary Figure 4: Mosaic white eye phenotype observed in transheterozygotes.** Left to right – wild-type, *kmo*-gRNA, *vasa*-Cas9, *vasa*-Cas9 + *kmo*-gRNA double heterozygote. **(A)** Images taken under white light. **(B)** Images taken under mCherry fluorescent filter. Mosaic white eye phenotype (disrupted eye pigments) can only be observed in double heterozygotes (white arrows), as discerned by fluorescence microscopy.

**SUPPLEMENTARY TABLES**

**Supplementary Table 1**

| **List of oligonucleotides used in the study.** | |
|---|---|
| **Name** | **Primer** |
| Primers used for building *w*6-gRNA-eGFP construct | |
| V1354 | GAGAAGAAATTTCGCTACTGTCAC |
| V1386 | ATCCCCTACCGGTCAGATTTCTC |
| V1390 | CTCATACTTGATTGTGTTTTACGCGAGAACACGTTGATCACGGCAAACAC |
| V1392 | GCAAGTCGTCGTCGTCGTCTGGGCCGCGGAGCTGCCGGTGTTTTTG |
| V0999 | CGCGTAAAACACAATCAAGTATGAG |
| V1259 | GCCTTTGAGTGAGCTGATACCATTGCAAGTTCCTAACCATAACCTAC |
| V1391 | GGCTGATTATGATCAGTCGACCAAGGAACACCTGTTCCCTCTG |
| V1362 | CCCAGACGACGACGACGACTTGC |
| V1406 | GCAGGAACACGGGCAATTCAGCAgAGAACACGTTGATCACGGCAAACAC |
| V1407 | GCTTGGATAGCGATTCGAGTTAACGCGCGTAAAACACAATCAAGTATGAG |
| V1405 | CTGCTGAATTGCCCGTGTTCCTGC |
| V1360 | CGTTAACTCGAATCGCTATCCAAGC |
| Primers used for genotyping of transformants | |
| V1476 | ATGGTTGCTGTGCACCCTTAACC |
| V1514 | GAGGTAATGCACGTATCCGCTT |
| V1477 | GTCGCTAGGTTAACCATGTGTAGG |
| V1515 | CTTCCCTCCGTTGTCTGGAG |

### SUPPLEMENTARY DATA

**Supplementary Data 1 - The *w*6-gRNA drive injection and transgenesis counting data**

This table contains the raw counting data of G1 with eGFP positive(GFP) phenotype indicating the transgenesis occurrence. All hatched G0s were divided into male and female pools and crossed with wild-type individuals. Egg rafts from each pool were hatched and counted together.

**Supplementary Data 2 - The establishment of *w*6-gRNA transgenic line**

The recovered eGFP+/DsRed transgenic male was mated to wild-type females to establish a transgenic line. The G1s were divided and scored by different sex and fluorescent markers, and the data is reported in the table. The inheritance ratio of the w6-gRNA drive element was calculated by dividing the number of individuals carrying eGFP positive (GFP) by the total number of G1s.

**Supplementary Data 3 - The *white* locus GD data**

Raw counting data of the G2 progeny with phenotypic scoring for DsRed positive (DsRed), eGFP positive (eGFP), *white-*/*white-*, or no fluorescence (none). The Cas9 transgene was tracked by DsRed presence and gRNA transgene was tracked by eGFP presence. Transgene inheritance rates in G2 for each single-pair cross (marked as "G1 (single-pair) cross" in the table) were calculated. Average inheritance and standard deviation were calculated for each transgene as well. The data is subdivided into the following tabs:

1. Fig. 2c - **The male copying data**: Counting data and inheritance rates with the *w*6-gRNA derived from G1 male germlines.
2. Fig. 2d - **The female copying data:** Counting data and inheritance rates with the *w*6-gRNA derived from G1 female germlines.
3. Fig. 3b,c - **Marked chromosome w4 homing data**: Counting data, average inheritance, and conversion rates with a marked chromosome approach.

**Supplementary Data 4 - The *kmo* locus GD data**

Raw counting data of the G2 progeny with phenotypic scoring for Cas9 only (*Opie2*-DsRed), Cas9 and *kmo*-gRNA (*Hr5IE1*-DsRed) together, *kmo*-gRNA (*Hr5IE1*-DsRed) only, or wildtype (no fluorescence). The Cas9 transgene was tracked by *Opie2*-DsRed presence and *kmo*-gRNA transgene was tracked by *Hr5IE1*-DsRed presence which can be phenotypically distinguished (Supplementary Figure 3). Transgene inheritance rates in G2 were calculated. Average inheritance and standard deviation were calculated for each transgene as well.
